# SNooPy: a statistical framework for long-read metagenomic variant calling

**DOI:** 10.64898/2025.12.01.691549

**Authors:** Roland Faure, Ulysse Faure, Tam Truong, Alessandro Derzelle, Dominique Lavenier, Jean-François Flot, Christopher Quince

**Affiliations:** Sequence Bioinformatics, Department of Computational Biology, Institut Pasteur, Paris, France; Université de Rennes, Inria, CNRS, IRISA - UMR 6074, Rennes, France; Service Evolution Biologique et Ecologie, Université libre de Bruxelles (ULB), Brussels, Belgium; Dep. of Mathematics, ETH Zürich, Switzerland; Organisms and Ecosystems, Earlham Institute, Norwich, UK; Gut Microbes and Health, Quadram Institute, Norwich, UK; School of Biological Sciences, University of East Anglia, Norwich, UK

## Abstract

Current long-read single-nucleotide variant callers were designed primarily for genomic data—particularly human genomes. While some have been used on metagenomic data, their underlying assumptions and training procedures fail to account for the inherent complexity of metagenomic samples. To date, no long-read variant caller has been purpose-built for metagenomic applications. To address this gap, we present SNooPy, a SNP-calling tool that implements a new statistical framework tailored to long-read metagenomic data. Unlike previous genomic methods, our approach makes no assumptions about the number of haplotypes present, their evolutionary relationships, or their sequence divergence. We demonstrate that SNooPy outperforms both traditional statistical and deep learning–based SNP callers. Our results suggest that future integration of this framework with deep learning approaches could further enhance variant calling performance. SNooPy is freely available on github.com/rolandfaure/snoopy.

## Introduction

A fundamental problem in the analysis of genomics data is the detection of variants: given a consensus genome from one individual and sequence reads from a related but not identical organism, how to compare the two. Differences can comprise single nucleotide variants (SNVs) or more complex structural variations such as rearrangements, insertions and deletions. Variant calling is the first step in numerous genomics applications, from the construction of genealogies to genome wide association studies (GWAS) [22, 25]. It is also a key step in metagenome analysis, where reads derive from multiple different organisms, typically microbes, and encompass not only inter-species diversity but also intra-species strain variation [23, 18]. Increasingly, metagenome analysis focuses on this strain-level, requiring variant identification within species [19, 16].

Not surprisingly given the importance of this problem, many programs for genomic variant calling have been developed. At present, two main families of variant callers exist. Historically, the first approach, statistical variant callers,employed probabilistic models to differentiate true genetic variants from sequencing errors [10, 11, 6, 7, 8]. These models were built upon assumptions regarding the data, typically that sequencing errors were independent or that the sequenced samples were diploid. A second paradigm emerged following the introduction of DeepVariant in 2016 [17], leveraging deep neural networks [14, 29, 1]. These replaced the statistical tests with “black-box” machine-learning, which requires training using ground-truth datasets of known variants. While these callers have achieved state-of-the-art performance for human genomes, their accuracy remains heavily dependent on the training data, sometimes underperforming when calling variants outside of the set of species on which they were trained [28].

In metagenomes, multiple strains of the same species may exist with very different abundances. The species genomes may also represent novel, previously unseen diversity, generated by binning de novo assemblies into metagenome-assembled genomes (MAGs) [19]. These specificities of metagenomes compared to single genomes can break assumptions behind some statistical models (typically the assumption that the sample is diploid or polyploid). Neural network callers will have been typically trained on known genomes (e.g. human) and are also implicitly learning the biases of their training data, which might be specific to the genome and not apply to metagenome use-case. Metagenomics hence calls for a specific approach to variant calling. Several short-read variant callers have been developed to address this challenge [2, 16, 6]. However, to the best of our knowledge, no long-read variant callers have been specifically developed for metagenomic applications.

To fill this methodological gap, we present SNooPy, a novel statistical SNP caller tailored for long-read metagenomic datasets. Similar to existing tools such as Longshot [8] and NanoCaller [1], SNooPy exploits the statistical dependencies among reads that arise from the inherent population structure of the sequenced sample. However, unlike Longshot and NanoCaller, SNooPy’s statistical framework is designed for metagenomic samples and does not make any assumption about ploidy. To do so, we implement a new statistical test inspired from previous work on haplotype assembly [9], which makes no assumptions on the number of haplotypes, their sequencing depth, or the sequencing error profiles: its only assumption is that sequencing errors occur independently across distinct reads.

Because of the lack of existing specialized metagenomic long-read SNP callers, we chose to compare SNooPy (0.4.3) with the widely used genomic SNP callers bcftools (1.22), Longshot (1.0.0), Nanocaller (3.6.2), Clair3 (1.0.10), and Deepvariant (1.9.0). These were therefore run slightly outside of their intended application area but it is common in metagenomics applications to use genomic variant callers. As detailed below, this demonstrated that SNooPy significantly outperformed not only statistical methods but also deep-learning ones (when applied without retraining, as is usually the case), and, hence, provides a fast and effective means to detect variants even in noisy long-read metagenome data.

## MATERIALS AND METHODS

### Multi-loci variant calling

SNooPy employs a multi-locus analysis strategy, i.e., it does not attempt to detect individual variants but rather identify and validates variant groups. This approach leverages the statistical principle that sequencing and alignment errors occur (nearly) independently; therefore, the probability of observing correlated sequencing errors across multiple reads at numerous loci is vanishingly small. When correlated patterns are detected, they more likely indicate reads originating from a distinct strain carrying true variants, rather than sequencing artifacts.

This work builds upon ideas developed in the field of metagenome assembly, more specifically upon the software HairSplitter [9] and is based on the code of Strainminer [24]. This section focuses on the core variant-calling procedure, which comprises two primary steps: (1) identification of correlated loci representing candidate groups of single nucleotide polymorphisms (SNPs), and (2) statistical validation of these candidate groups. The complete variant-calling algorithm includes additional validation steps and recovery mechanisms, detailed in subsection *Implementation details*.

SNooPy processes BAM files through a sliding window approach. Within each window, the pileup data is transformed into a binary matrix *M*, where rows represent individual reads, columns correspond to genomic loci, and each entry *M*_*ij*_ equals 1 if read *i* contains the reference allele at locus *j*, or 0 if it contains an alternative base. To identify candidate variants, the algorithm starts by computing pairwise correlations between all columns using chi-square tests, yielding a p-value for each pair of columns. This p-value is used to perform complete-linkage hierarchical clustering with a p-value cutoff of 0.05 to group highly correlated columns. The choice of complete-linkage clustering ensures that all pairs of columns correlate well. From these groups, we identify variant patterns as subsets of loci and reads where all bases show a non-reference allele.

Building upon the statistical framework established in HairSplitter [9], we developed a statistical test which evaluates whether observed variant patterns represent authentic variants (alternative hypothesis) or result purely from coincidental sequencing errors (null hypothesis). When a pattern passes the test, the corresponding variants are outputted in a VCF file.

Let *a* be the number of reads and *b* the number of loci in the pattern, among a matrix totalling *n* reads and *m* loci. Let *s* denote an upper bound on the per-base sequencing error probability, estimated from the alignment data as described below.

In a matrix of size *n* × *m*, there are 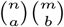 submatrices of size *a* × *b*. In any of these submatrix, under our independence assumption, the probability that *all* the bases of the submatrix are sequencing errors is ≤ *s*^*ab*^. The union bound gives us a bound on the probability *p* of observing such a pattern *at least once* among the 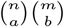 submatrices under our null hypothesis: 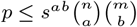 . We reject the null hypothesis when *p* ≤ 0.001.

To enhance computational efficiency and statistical power, we only include loci where alternative alleles appear in more than 5% of reads. This filtering substantially reduces the matrix dimensions, accelerating computations while strengthening the statistical test through a reduced value of *m*. The error rate is estimated as the divergence between the reads and the reference. To account for error-prone regions such as homopolymers and ensure that our error rate parameter *s* always remains higher than the actual local error rate, we empirically set *s* as three time the measured error rate.

For illustrating the statistical strength of the procedure, consider a pileup of 100 Oxford Nanopore reads with error rate 0.05 (hence *s* = 3 × 0.05 = 0.15) spanning 5,000 base pairs. Let us imagine that among these 5,000 loci, 500 exhibit an alternative allele in more than 5% of reads and that we observe 5 reads sharing alternative bases at 10 loci. The probability that this pattern arises from sequencing errors alone is ≤ 3 *×* 10^−17^, providing strong evidence for genuine variants. A graphical illustration of the test is shown on Figure 1.

**Fig. 1:**
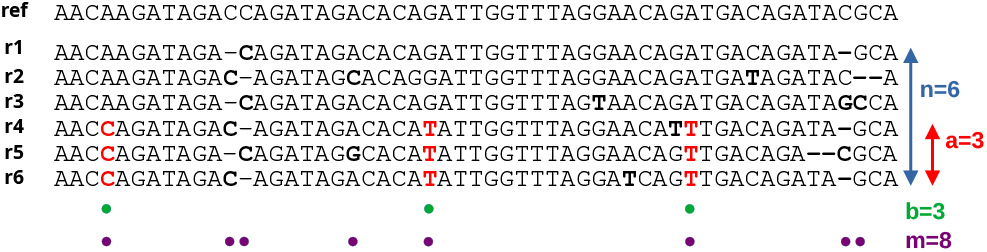
Statistical foundation of the SNooPy algorithm. SNooPy starts by identifying groups of correlating columns, highlighted by green circles (*b*), among all columns potentially containing variants, highlighted by purple circles (*m*). *n* is the total number of reads and *a* the number of reads bearing the tested variants. The error rate *s* is over-estimated as three times the divergence of the reads to the reference. Here there are 31 non-reference bases out of 336, hence *s* = 3 × 31*/*336 = 0.28. Our statistical test states that the probability of observing this pattern due to independent sequencing errors is 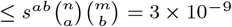.

### Implementation details

#### Rescuing SNPs

Our multi-locus variant calling approach has three potential limitations that could result in missed SNPs. We have developed specific rescue strategies to address each limitation.

#### Detection of isolated SNPs

Multi-locus variant calling relies on correlations between SNPs to achieve statistical power. However, isolated SNPs lacking correlation with other variants cannot benefit from this approach. While our statistical test remains valid for single SNPs, its power is substantially reduced.

To rescue isolated SNPs, we implement a position-specific analysis using a simple binomial model. For each genomic position, we model the expected number of sequencing errors as following a binomial distribution with parameters *c* (the coverage at that position) and *s* (the maximum error rate). When an alternative allele appears at a frequency that yields a p-value smaller than 0.001 under this null model, we classify the position as an “obvious” SNP, bypassing the need for multi-locus validation.

#### Recovery of high-noise correlated variants

Some true SNPs may be difficult to detect due to elevated local error rates. During initial hierarchical clustering, such positions are excluded from variant groups to preserve the statistical power of our multi-locus test, as including high-noise positions can weaken the statistical test by excluding rows containing errors from the tested submatrix.

After establishing high-confidence variant calls, we perform a recovery phase where all loci are tested for correlation with confirmed SNPs using chi-square tests. Positions demonstrating significant correlation (p-value *<* 10^−6^) with established variants are rescued and called as SNPs.

#### Detection of multi-allelic sites

At positions harboring multiple alternative alleles, our primary algorithm typically identifies only the most frequent variant, potentially masking additional polymorphisms. To address this limitation, we perform iterative variant calling. After the initial round, all called variants are masked (i.e., converted to reference-like status in the binary matrix), and the algorithm is re-executed on the modified data. This iterative process reveals previously hidden alternative alleles by removing the dominant variant signal, allowing detection of secondary polymorphisms at multi-allelic sites.

#### Transforming a read alignment into matrices

The algorithm begins by transforming read alignments from BAM format into binary matrices suitable for variant calling. Only the best alignment of each read is retained. Computing all-versus-all locus correlations scales quadratically, making it extremely computationally expensive on long contigs. However, two loci can only exhibit meaningful correlation if they are close enough to be spanned by a significant number of reads. Therefore, we lose minimal information by restricting correlation calculations to nearby loci. To implement this, we partition the reference genome into fixed-length windows, setting each window to half the median read length.

Read mapping patterns provide additional information for strain identification. Reads frequently map to only partial segments of the reference genome rather than aligning end-to-end, especially when they come from strains with structural variations regarding the reference. To exploit this signal, we cluster reads within each window based on their mapping coordinates. Groups of at least 5 reads sharing exactly the same mapping coordinates are grouped together, as they potentially originate from the same strain. Conversely, reads mapping to different coordinates within the window are analyzed separately, as they probably originate from distinct strains with different genomic architectures or insertion/deletion patterns. This coordinate-based clustering serves as a pre-filtering step that segregates reads before variant calling. By analyzing each read cluster independently, we prevent the conflation of signals from different strains and enhance the algorithm’s ability to detect strain-specific variants. This approach is particularly effective when strains exhibit structural variations that cause their reads to map to different reference coordinates within the same genomic window.

## RESULTS

### Benchmark description

We compared SNooPy (0.4.3) with the widely used SNP callers bcftools (1.22), longshot (1.0.0), Nanocaller (3.6.2), Deepvariant (1.9.0) and Clair3 (1.0.10), all run with recommended or default options.

We benchmarked SNooPy on three sequencing datasets. The first one is a commercially available mock community, named Zymobiomics Gut Microbiome Standard, sequenced using ONT R10.4.1 (SRR17913199). This community has the particularity of containing five strains of *Escherichia coli* which are mostly collapsed in the metaFlye assembly, in which we expect to observe variants. The second and third ones are a human stool and a soil sample sequenced in [4] using the latest ONT R10.4.1 flow cells. Both of these datasets were chosen because PacBio HiFi sequencing of the same samples were conducted [3] and were thus available to evaluate the quality of the calls .

All datasets were assembled using metaFlye [12]. We then called the variants using the assembly as a reference. For the Zymobiomics Gut Microbiome Standard, we report recall and precision only on the *E. coli* contigs, to measure the ability of the SNP callers to call variants in a multi-strains context. The soil dataset presented computational challenges due to its very large size (6.8G), which exceeded the processing capacity of all tools within our one-week runtime constraint. To address this limitation, we randomly selected 921 contigs (with an N50 of 56kb) and conducted our analysis exclusively on this subset.

We encountered a technical issue with DeepVariant, which crashes when processing loci containing multiple alternative alleles. We have reported this bug to the DeepVariant GitHub repository. As an interim solution, we excluded the problematic contigs from our analyses. Since this exclusion did not significantly alter the performance metrics of other variant callers, we report statistics across the restricted dataset for all tools.

To confirm our analyses, we created simulated datasets, and benchmarked the variant callers on them. More precisely, we selected 10 *E. coli* genomes spread across the phylogenetic tree of *E. coli*. We then simulated Nanopore sequencing using meta-NanoSim [27], trained to mimic the error pattern of dataset SRR17913199, varying the number of strains and their coverage. We also simulated Nanopore sequencing using Badread [26] using the “nanopore2023” model, which allowed us to vary the error rate. The experiments are detailed in Supplementary Table 1.

### Evaluation metrics

We assessed the recall and precision of the SNP callers. Variant call comparison is challenging because SNP callers implicitly assume that reads align on the reference end-to-end, an assumption that fails with highly divergent sequences or structural variants. We therefore excluded variants longer than 5bp from our analysis (less than 0.2% of the variants), as different SNP callers may legitimately disagree on these calls. For transparency and reproducibility, all comparison scripts used in this analysis are available in our GitHub repository (github.com/RolandFaure/SNooPy).

For the Zymobiomics dataset, we aligned the reference genomes against the assembly and used this alignment to build a set of ground truth SNPs.

For the other two datasets, we employed PacBio HiFi sequencing reads, which we mapped on the assemblies using minimap2 [13], and validated or invalidated variants based on their presence in the alignment of the set of HiFi reads on the same assemblies. This approach has inherent limitations: namely, the two sequencing experiments might have only sequenced a (different) sample of the overall diversity. Nevertheless, the detection of a variant using both technologies strongly supports its validity. For these two datasets, we defined recall as the proportion of HiFi-confirmed SNPs that each software successfully identified, relative to the total pool of HiFi-confirmed SNPs called by any software. This definition intentionally excludes variants observed exclusively in HiFi data. The precision metric requires careful interpretation in this context. When SNPs called from ONT data lack confirmation in HiFi data, we classify them as false positives. However, this classification may be overly stringent: some of these variants may represent true polymorphisms that simply were not captured in the HiFi sequencing. Indeed, manual investigation using Logan-Search [5] against SRA revealed that several supposed “false positives” had been previously observed in other datasets, suggesting they are likely genuine variants. The limited throughput of Logan-Search did not allow us to conduct a systematic analysis of all these putative false positives.

The scripts to normalize, merge and compare the obtained VCFs to either the ground truth or the HiFi mapping results are available at https://github.com/rolandfaure/snoopy.

### SNooPy excels on complex communities

Figures 2 and 3 present the recall and precision metrics obtained on real and simulated datasets, respectively (exhaustive results are provided in the supplementary data). SNooPy consistently achieved the highest recall across all datasets, demonstrating superior performance for metagenomic variant calling

**Fig. 2:**
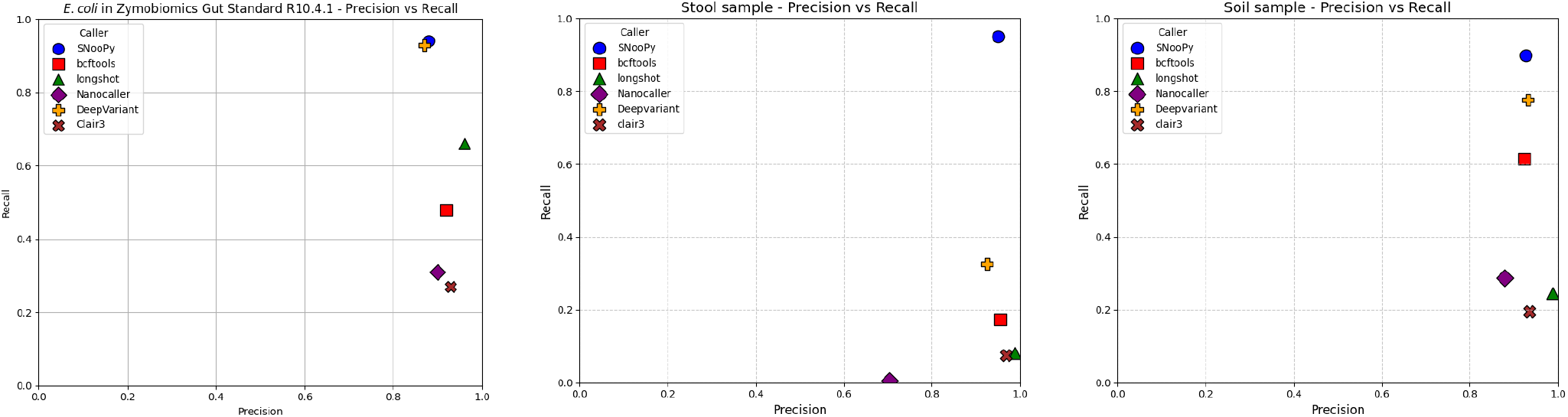
Precision and recall of benchmarked metagenomic tools on three datasets. **Left:** ONT sequencing of a synthetic gut microbiome mock community. Metrics are calculated only against known *E. coli* genomes. **Middle and right:** ONT R10.4.1 sequencing of a human stool sample and a soil sample. Metrics are evaluated using a PacBio HiFi sequencing of the same sample. Recall is computed w.r.t. the union of all called variants confirmed by HiFi, as some variants may be present exlusively in the HiFi sequencing.

**Fig. 3:**
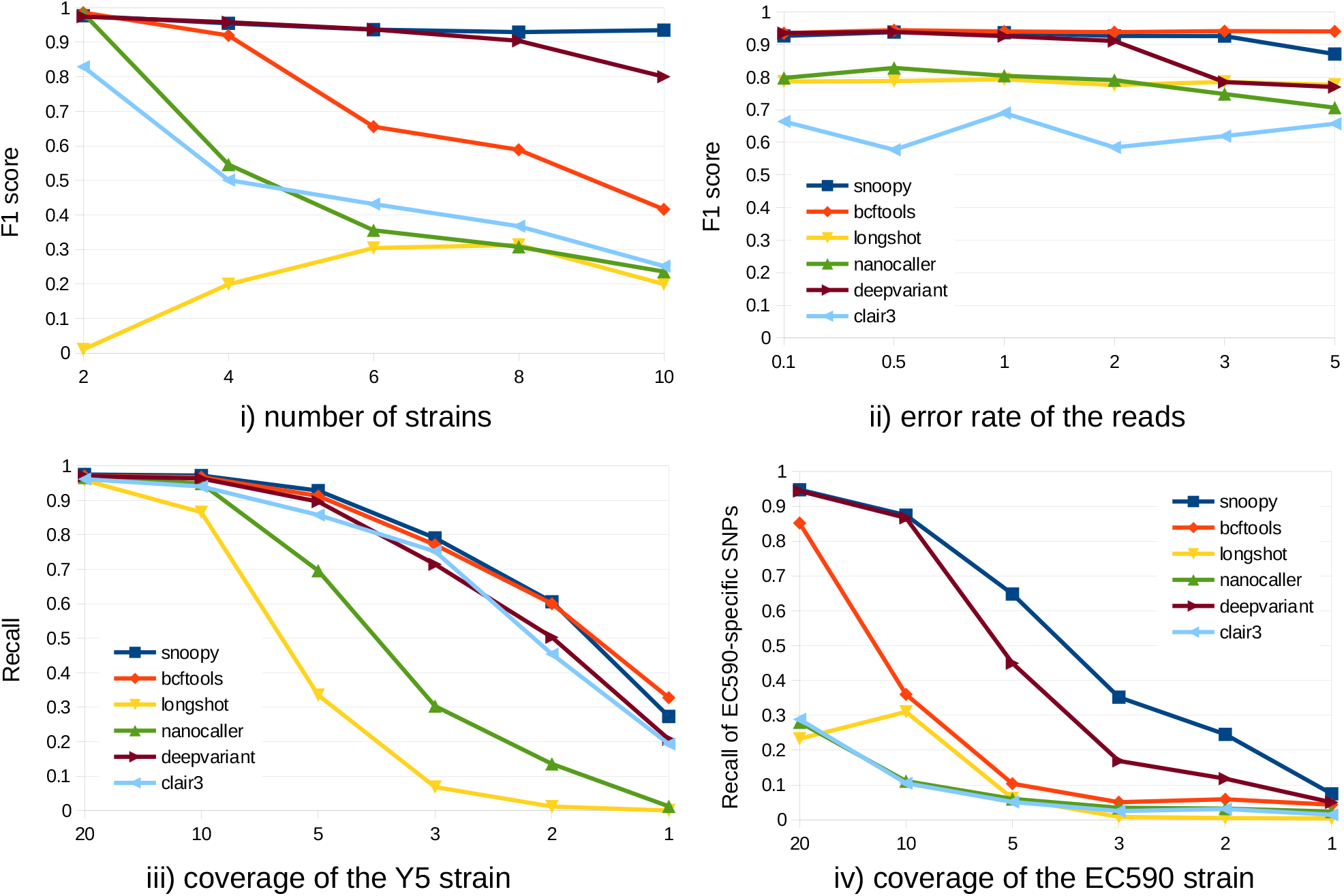
Recall or F1 metric (i.e. harmonic mean of precision and recall) of variant callers evaluated on simulated read datasets with varied parameters: (i) Mixtures of two to ten strains, with 20× coverage for all strains. Variants called against metaFlye assembly. (ii) Four-strain mixtures, reads generated with Badreads with varying error rates and 20× coverage for all strains; variants called against metaFlye assembly; (iii) Y5 strain sequenced at different coverages; variant called against H5 genome; (iv) Four-strain mixture, three strains at 20× coverage, and EC590 at varying coverage. Variants called against metaFlye assembly. For (iv), only recall of EC590-specific SNPs is reported.

Clair3, Nanocaller, Longshot and, to a lesser extent, Bcftools, showed limited recall in metagenomic contexts, in line with their documentation that specifically targets diploid applications. Their performance degraded dramatically when the number of strains increased (experiment (i)) or when the coverage became uneven between the strains (experiment (iv)).

DeepVariant matched the performance of SNooPy on the *E. coli* strains of the mock community, achieving more than 90% recall, but called 13% fewer variants than SNooPy on the soil sample and detected barely a third of all variants detected by SNooPy on the human gut sample. The simulated experiments elucidate these differences: DeepVariant matches SNooPy’s performance up to six equally-abundant strains (experiment (i)), equivalent to the mock community configuration. However, DeepVariant shows reduced recall under low-coverage conditions (experiments (iii) and (iv)), the dominant scenario in the soil sample (11× average coverage). SNooPy also exhibit superior performance in strain-rich environments and with unequal strain coverage (experiments (i) and (iv))—conditions characteristic of the human gut sample (82× average coverage).

We analysed variants detected by DeepVariant but missed by SNooPy. The vast majority of missed variants exhibited very low coverage (≤4× for 90% of cases), as shown in Supplementary Figure 1. Manual inspection of a subset of variants revealed that DeepVariant can outperform SNooPy in high-noise alignment regions, for example around homopolymeric stretches. We attribute this advantage to the deep neural network’s capacity to learn robust features in regions where sequence alignment is difficult. Despite this, SNooPy’s overall recall remained superior to DeepVariant, even when considering only low-coverage variants.

SNooPy processed the Zymobiomics, gut and soil samples in 20 minutes, 2h30 and 1h on 48 threads, respectively. This is comparable to DeepVariant, which completed in approximately one hour for all the three datasets. The complete list of running times and RAM usage is available in supplementary data.

## DISCUSSION

In this study, we present a new approach for long-read metagenomic variant calling based on a simple, non-parametric test of correlation among reads. To our knowledge, this represents the first statistical variant-calling framework for long reads built on assumptions sufficiently general to hold across virtually all types of sequencing experiments, including metagenomic data. We implemented this test in SNooPy, a metagenomic variant caller that performed best among the methods tested, including the deep-learning state-of-the-art methods. This benchmark shed light on the limitations of the majority of existing long-read variant callers when applied to metagenomics data. This was unsurprising given the data they were trained on and their underlying assumptions, but represent an important, under-appreciated caveat for practitioners developing their own long-read metagenomics analysis pipelines. For instance, Longshot has been frequently used in metagenomics contexts [21, 20] but performed poorly in our tests. DeepVariant stands out as an exception, providing adequate precision and recall in the metagenomic context. However, the current release (v1.9.0) suffers from a bug that hampers its practical application in metagenomic analyses.

The strength of our pileup-based statistical variant calling is that it is grounded in a solid statistical framework explicitly designed for metagenomics. By contrast, the strength of deep neural networks, which take whole alignments as input rather than considering only pileups at few distinct loci, lies in its ability to draw information even out of noisy, low-coverage alignments [14]. We believe that both approaches are in essence orthogonal and could be combined to exploit the strengths of both strategies. For instance, the information of co-occurring variants used in our statistical test could be incorporated as an input feature of a deep learning variant caller. The result could be a method developed for long-read metagenomics that would improve the effectiveness of SNooPy in regions with ambiguously aligned reads.

Our benchmark on the soil sequencing dataset shows that even the best-performing tool, SNooPy, missed 10% of the SNPs. Furthermore, this is a lower estimate as our metric did not account for SNPs missed by all callers. Given the increasing importance of both long-read sequencing and strain analysis in metagenomics, and the potential for improvement that this indicates, the development of dedicated long-read metagenomic variant callers is likely to remain an active research field in the coming years.

## Supporting information

exhaustive results are provided in the supplementary data

## Software availability statement

SNooPy is freely available on github.com/rolandfaure/snoopy under a MIT licence.

## Data availability statement

SNooPy is freely available at https://github.com/rolandfaure/ SNooPy and on Zenodo with DOI 10.5281/zenodo.19205710. The sequencing datasets are freely available with accession numbers ERR15285694 (human gut ONT), ERR15289675 (human gut HiFi), ERR15289757 (soil ONT), ERR15289804 (soil HiFi) and SRR17913199 (Zymobiomics Gut Microbiome Standard ONT). Zymo mock reference genomes are available at https://s3.amazonaws.com/zymo-files/BioPool/D6331.refseq.zip. All reference genomes and command lines used to run the benchmark are available on Zenodo, 10.5281/zenodo.19218967.

## Supplementary Data Statement

Supplementary Data are available at NAR Online.

## Author Contributions Statement

**Roland Faure:** Investigation, Conceptualization, Software, Writing. **Ulysse Faure:** Conceptualization. **Tam Truong:** Software. **Alessandro Derzelle:** Investigation. **Dominique Lavenier:** Conceptualization, Supervision. **Jean-François Flot:** Conceptualization, Supervision. **Christopher Quince:** Conceptualization, Supervision, Writing.

## Funding

R.F. is supported by the Horizon Europe ERC grant number 101088572 “IndexThePlanet”. C.Q. acknowledges the support of the Biotechnology and Biological Sciences Research Council (BBSRC), part of UK Research and Innovation; Earlham Institute Strategic Programme Grant (Decoding Biodiversity) BBX011089/1 and its constituent work package BBS/E/ER/230002C; the Core Strategic Programme Grant (Genomes to Food Security) BB/CSP1720/1 and its constituent work packages BBS/E/T/000PR9818 and BBS/E/T/000PR9817; and the Core Capability Grant BB/CCG2220/1.

## ACKNOWLEDGEMENTS

We acknowledge the GenOuest bioinformatics core facility (https://www.genouest.org) for providing the computing infrastructure. Many thanks to Rumen Andonov for his feedback, brainstorming and careful proofreading. The program Tablet [15] was used to visualize data while developing SNooPy. Various LLM have been used to improve the writing. For the purpose of Open Access, a CC-BY public copyright licence has been applied by the authors to the present document and will be applied to all subsequent versions up to the Author Accepted Manuscript arising from this submission.

## Conflict of interest statement

None declared.

## Notes

https://www.doi.org/10.5281/zenodo.19218966

### Competing Interest Statement

The authors have declared no competing interest.

### Summary of Updates

The software has been updated, leading to new results.

