## Supplementary Table 1 for "SNooPy: a statistical framework for long-read metagenomic variant calling"

April 2, 2026

| | | strains | coverage | read simulator | reference |
|---|---|---|---|---|---|
| Number of strains | 2 | Y5 H5 | 20x | NanoSim<br>ONT R10.4.1 | Flye assembly |
| | 4 | Y5 H5<br>AMSCJX03 EC590 | 20x | NanoSim<br>ONT R10.4.1 | Flye assembly |
| | 6 | Y5 H5<br>AMSCJX03 EC590<br>K12 LD27-1 | 20x | NanoSim<br>ONT R10.4.1 | Flye assembly |
| | 8 | Y5 H5<br>AMSCJX03 EC590<br>K12 LD27-1<br>ME8067 RM14721 | 20x | NanoSim<br>ONT R10.4.1 | Flye assembly |
| | 10 | Y5 H5<br>AMSCJX03 EC590<br>K12 LD27-1<br>ME8067 RM14721<br>SE15 UMN026 | 20x | NanoSim<br>ONT R10.4.1 | Flye assembly |
| Error rate (%) | 0.1 | Y5 H5 AMSCJX03 EC590 | 20x | BadReads<br>ONT2023, 0.1% errors | Flye assembly |
| | 0.5 | Y5 H5 AMSCJX03 EC590 | 20x | BadReads<br>ONT2023, 0.5% errors | Flye assembly |
| | 1 | Y5 H5 AMSCJX03 EC590 | 20x | BadReads<br>ONT2023, 1% errors | Flye assembly |
| | 2 | Y5 H5 AMSCJX03 EC590 | 20x | BadReads<br>ONT2023, 2% errors | Flye assembly |
| | 3 | Y5 H5 AMSCJX03 EC590 | 20x | BadReads<br>ONT2023, 3% errors | Flye assembly |
| | 5 | Y5 H5 AMSCJX03 EC590 | 20x | BadReads<br>ONT2023, 5% errors | Flye assembly |
| Even coverage | 20 | Y5 | 20x | NanoSim<br>ONT R10.4.1 | H5 |
| | 10 | Y5 | 10x | NanoSim<br>ONT R10.4.1 | H5 |
| | 5 | Y5 | 5x | NanoSim<br>ONT R10.4.1 | H5 |
| | 3 | Y5 | 3x | NanoSim<br>ONT R10.4.1 | H5 |
| | 2 | Y5 | 2x | NanoSim<br>ONT R10.4.1 | H5 |
| | 1 | Y5 | 1x | NanoSim<br>ONT R10.4.1 | H5 |
| Uneven coverage | 20 | Y5 H5 AMSCJX03 EC590 | 20x, 20x, 20x, 20x | NanoSim<br>ONT R10.4.1 | Flye assembly |
| | 10 | Y5 H5 AMSCJX03 EC590 | 20x, 20x, 20x, 10x | NanoSim<br>ONT R10.4.1 | Flye assembly |
| | 5 | Y5 H5 AMSCJX03 EC590 | 20x, 20x, 20x, 5x | NanoSim<br>ONT R10.4.1 | Flye assembly |
| | 3 | Y5 H5 AMSCJX03 EC590 | 20x, 20x, 20x, 3x | NanoSim<br>ONT R10.4.1 | Flye assembly |
| | 2 | Y5 H5 AMSCJX03 EC590 | 20x, 20x, 20x, 2x | NanoSim<br>ONT R10.4.1 | Flye assembly |
| | 1 | Y5 H5 AMSCJX03 EC590 | 20x, 20x, 20x, 1x | NanoSim<br>ONT R10.4.1 | Flye assembly |

Supplementary Table 1: Description of the experiments run with the simulated datasets on *E. coli*

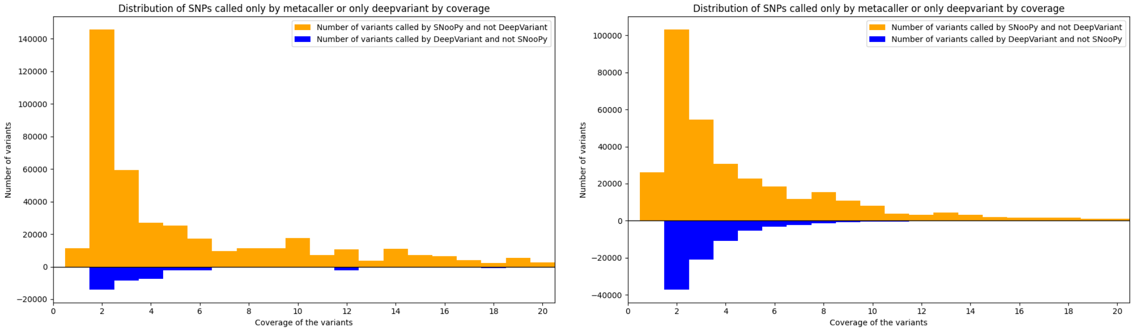


Supplementary Figure 1: Coverage of the variants missed by DeepVariant but not SNooPy, and inversely.
